# Microwear variability in a spatially and temporally constrained elephant population: Implications for interpreting the diets of extant and extinct proboscideans

**DOI:** 10.1101/2025.09.14.676112

**Authors:** Chase Alexander Barrett, Melissa Pardi, Larisa DeSantis

**Affiliations:** Department of Biological Sciences, Vanderbilt University, Nashville, TN, USA; Research & Collections Center, Illinois State Museum, Springfield, IL, USA; Department of Earth & Environmental Sciences, Vanderbilt University, Nashville, TN, USA; Evolutionary Studies, Vanderbilt University, Nashville, TN, USA

## Abstract

Proboscideans, including mammoths, played a crucial role in past herbivore communities, where resource partitioning helped reduce competition and promote coexistence. Stable carbon isotopes are frequently employed to differentiate between the consumption of C_3_ and C_4_ plants in the fossil record. However, as geographic variability influences δ^13^C values, dental microwear texture analysis (DMTA) is often used in tandem to infer dietary preferences among extinct taxa (e.g., the consumption of grass vs. browse). Interpreting the dietary ecology of proboscideans, like mammoths, rests on our ability to compare fossil specimens to modern taxa with known diets. Here, we established a modern reference for interpreting mammoth DMTA by analyzing teeth from 11 African bush elephants (*Loxodonta africana*) culled in 1993 from Kruger National Park (KNP), South Africa (CITES permit certificate no. 780873). These specimens, housed at the Illinois State Museum, originated from the arid bushveld of northern KNP and were collected during the beginning of the dry season. Previous studies indicate that modern elephants in this region consume a mixed diet, consisting of ∼40% grass in the dry season and 50% grass in the wet season. The well-documented dietary and environmental context of these individuals provides an opportunity to assess dental microwear patterns in a modern analog and compare individuals with known diets to extinct mammoths. Specifically, we compared newly acquired modern African bush elephants DMTA to published DMTA data from fossil specimens of Columbian mammoths (*Mammuthus columbi*). Comparisons with fossil assemblages reveal few statistically significant differences in microwear between mammoth and KNP elephants, with the exception of mammoths from Leisey Shell Pit 1A having significantly lower complexity values than modern African bush elephants—indicative of some mammoth populations eating softer foods and/or less woody browse. Variation and breadth of DMTA from mammoths are similar to the temporally and geographically constrained population of *L. africana*. Despite potential time averaging in fossil assemblages, the variation in mammoth DMTA aligns with that of a geographically and temporally constrained modern population, indicating that microwear variability in fossil taxa is not necessarily greater than that observed in extant species and is consistent with the highly varied diets of modern African elephants.

## Introduction

Proboscideans, including extinct forms such as mammoths (*Mammuthus*) and mastodons (*Mammut*) and extant species like African (*Loxodonta*) and Asian (*Elephas*) elephants, have long been keystone members of herbivore communities (Abraham et al. 2024). As megaherbivores, they play critical roles in shaping vegetation structure through disturbance, influencing nutrient cycling, and facilitating seed dispersal across ecosystems (Owen-Smith, 1987; Aarde et al., 1999; Poulsen et al., 2018; Kamga et al., 2022). Fossil evidence suggests that proboscideans occupied dietary niches ranging from grazers to mixed feeders to browsers, depending on species, region, and climatic conditions (Feranec and Macfadden, 2000; Koch et al., 2004; Green et al., 2017; Smith et al., 2018; Pardi & DeSantis, 2022). Given their ecological significance, understanding their dietary variability is crucial for reconstructing past environments and assessing the mechanisms that facilitated proboscidean coexistence in diverse Pleistocene ecosystems.

Dental microwear has become an essential tool for reconstructing the diets of extinct mammals (Walker et al., 1978), complementing δ^13^C stable isotope analysis by providing high-resolution dietary data over shorter timeframes. Unlike isotopic proxies, which typically reflect long-term dietary averages, dental microwear captures feeding behavior in the days to weeks preceding death (Grine et al., 1988). Further, dental microwear tools have evolved since their inception and now focus on imaging dental microwear surface textures (i.e., dental microwear texture analysis, DMTA) in three dimensions via confocal microscopy and scale-sensitive fractal analysis (Ungar et al. 2003; Scott et al. 2005, 2006; DeSantis et al. 2013; DeSantis 2016). DMTA has since been extensively validated in modern taxa, including bovids (Scott et al., 2012), tapirs (DeSantis et al., 2020), and proboscideans (Smith & DeSantis, 2018; Smith & DeSantis, 2020), demonstrating the ability to differentiate between grazing, browsing, and mixed feeding strategies (DeSantis, 2016). In fossil proboscideans, DMTA has been used to interpret dietary variation across mammoths, mastodons, and gomphotheres, shedding light on ecological partitioning within these extinct groups (DeSantis, 2016; Green et al., 2017; Smith & DeSantis, 2020). This approach allows researchers to assess dietary plasticity, environmental adaptations, and potential competition between coexisting species.

There exists a concern in paleoecology that, within a given fossil locality, time averaging increases dietary variability observed through dietary proxies (Rivals et al., 2007; Kidwell and Tomasovych, 2013; Davis and Pineda-Munoz, 2016; Abraham et al., 2024). The reasoning behind this argument is that fossil assemblages often integrate individuals that lived in a given location across centuries to millennia (Behrensmeyer, 1982; Kidwell and Tomasovych, 2013; Fisher, 2018). It is hypothesized that localized dietary reconstructions may, therefore, reflect artificially inflated dietary breadth patterns rather than the actual ecological variability observed in a single population (Kidwell, 2013; Davis and Pineda-Munoz, 2016; DeSantis, 2016; Abraham et al., 2024). However, this hypothesis has not been rigorously tested, and the extent to which time averaging inflates dietary variability remains uncertain.

Here, we examined the dental microwear textures of *Loxodonta africana* from Kruger National Park (KNP) to test whether potentially time-averaged mammoth assemblages exhibit greater dietary variability than time-constrained modern elephant populations. Given the structural similarities of their teeth to mammoths, and the similarities in their inferred diets through other proxies (Ungar et al., 2003; Codron et al. 2006; DeSantis, 2016; Green et al., 2017), we use African bush elephants as a modern analog for extinct mammoth populations, allowing for direct comparison of dietary variation within a well-documented, temporally and environmentally constrained population. If mammoths show significantly greater variability in DMTA attributes than KNP elephants, it would support the idea that time averaging inflates dietary breadth observed in fossil proboscidean assemblages. Alternatively, if variation in mammoth DMTA is comparable to that of modern elephants, then variation observed in fossil populations may be ecologically meaningful, and it should not be assumed that time averaging is the primary driver of the dietary variability in mammoth populations without other supporting evidence. By using a living species with known dietary habits as a reference, this study provides insight into dietary variability and ecological plasticity of extinct proboscidean populations.

## Materials & Methods

Our modern sample consists of teeth from 11 individuals of *Loxodonta africana* that were culled as part of routine management activities in Kruger National Park (KNP) in 1993. These individuals are curated at the Illinois State Museum and were legally obtained under CITES permit certificate no. 780873. These individuals originated from the arid bushveld of the northern region of KNP in April/May (Aarde et al. 1999). Given the timing of the cull, the microwear from these animals reflects the diet consumed at the onset of the dry season. This is a period when elephants in the Northern region of KNP consume an estimated 40% grass as part of a mixed-feeding strategy (Codron et al. 2006). This dataset, therefore, provides a well-documented and spatially/temporally constrained sample for comparison with fossil mammoth populations. These data were subsequently compared to previously reported mammoth DMTA values from the following sites: Ingleside, Cypress Creek, and Friesenhahn Cave, TX, and Punta Gorda, Tri-Britton, and Leisey Shell pit 1A, FL (Smith & DeSantis, 2018; Smith & DeSantis, 2020). These sites are ideal for comparison, as they are well studied and the focus of both DMTA and stable isotope analyses on fossil proboscideans (Feranec & MacFadden 2000; Koch et al. 2004; Yann & DeSantis 2014; Smith & DeSantis, 2018; Lundelius et al., 2019).

Elephant teeth with occlusal wear were molded using polyvinylsiloxane dental impression material (President’s Jet regular body, Coltène-Whaledent Corp., Cuyahoga Falls, OH, USA). After casting the molds with Epotek 301 epoxy resin and hardener (Epoxy Technologies Corp., Billerica, MA, USA) and drying them for at least 72 hours, high-magnification microscopy was followed by scale-sensitive fractal analysis (SSFA) (Jones and DeSantis, 2016). Occlusal surfaces were scanned in three dimensions in four spatial quadrants (each individually measuring 102 × 138 µm^2^) using a 100x objective on a Sensofar Plu neox optical profiler at Vanderbilt University’s Department of Biological Sciences with a 0.73 numerical aperture, a sampling resolution of 36.33 data points per µm^2^, and a step height of 0.2 µm. The software Tooth Frax and SFrax (Surfract Corporation), were used to quantify surface attributes via scale-sensitive fractal analysis (Ungar et al., 2003; Scott et al., 2005, 2006). Anisotropy (exact-proportion length-scale anisotropy relief, *epLsar*) and complexity (area-scale fractal complexity, *Asf*c) provide quantitative measures of the similarity in orientation of wear features and the degree of surface texture variation across the occlusal surface, respectively. High *epLsar* in herbivores often suggests repetitive, uniform jaw movements typically associated with the consumption of tough, fibrous foods like grass and leaves (Ungar et al., 2003; Scott et al., 2005; Prideaux et al., 2009; Scott et al., 2012; Haupt et al., 2013; Jones & DeSantis, 2016; Hedberg & DeSantis, 2016; Jones & DeSantis, 2017; DeSantis et al., 2019). High *Asfc* reflects the roughness of the wear pattern, being characterized by more intricate and variable surfaces caused by the consumption of hard and brittle foods like woody browser, seeds, or nuts (Scott et al., 2005; Scott et al., 2012; DeSantis et al., 2015; DeSantis et al., 2019; DeSantis et al., 2020). Textural fill volume (*Tfv*) quantifies the volume of space that exists between peaks and valleys on a tooth’s worn surface by measuring the amount of material required to fill these features at defined scales. Higher *Tfv* values generally indicate coarser surface textures, which are often associated with the consumption of harder or more brittle foods that create deeper and more pronounced wear features (Scott et al., 2006; DeSantis, 2016).

Data collected from this sample (Table 1) were then compared to mammoth DMTA from previously published studies (Smith & DeSantis, 2018; Smith & DeSantis, 2020; Table 2). We used a Kruskal-Wallis test (non-parametric analysis of variance) to determine whether or not there were statistically significant differences within each DMTA parameter across our samples. We then used a Dunn’s test and Levine test to perform multiple comparisons to identify which groups were significantly different from the modern elephant population, via mean values and variance (standard deviation, n-1; Table 2), respectively. If time averaging in fossil assemblages artificially inflates variation in DMTA parameters, we would expect greater variation in fossil assemblages than in modern populations that are constrained by space and time.

**Table 1:** Kruger National Park dental microwear texture data. A table containing and comparing DMTA data sampled and collected from the *L. africana* specimen from Kruger National Park. Along with individual values, these data include the mean, medians, and standard deviation for each statistic.

| Catalog Number | *Asfc* | *epLsar* | *Tfv* | *HAsfc*<sub>3x3</sub> | *HAsfc*<sub>9x9</sub> |
| --- | --- | --- | --- | --- | --- |
| 154 | 2.806 | 0.00387 | 12907.64 | 1.9047 | 2.5962 |
| 55 | 4.539 | 0.00394 | 14811.06 | 0.9306 | 1.0348 |
| 159 | 1.842 | 0.00443 | 11803.94 | 0.1831 | 0.4329 |
| 163 | 5.336 | 0.00434 | 14348.65 | 0.5199 | 1.0409 |
| 189 | 3.916 | 0.00280 | 12231.40 | 0.1328 | 0.3268 |
| 536 | 5.809 | 0.00365 | 15732.41 | 0.1871 | 0.3879 |
| 538 | 2.286 | 0.00663 | 13836.97 | 0.3919 | 0.6476 |
| 63 | 7.829 | 0.00166 | 11803.94 | 0.1831 | 0.4329 |
| 64 | 3.122 | 0.00527 | 17384.12 | 0.2567 | 0.4660 |
| 65 | 1.299 | 0.00244 | 13673.70 | 0.5255 | 0.7503 |
| 66 | 3.397 | 0.00275 | 7928.31 | 0.1411 | 0.3067 |
| **Mean** | 3.835 | 0.00380 | 13314.74 | 0.4869 | 0.7657 |
| **Median** | 3.397 | 0.00387 | 13673.70 | 0.2568 | 0.4660 |
| **SD (n-1)** | 1.928 | 0.00139 | 2470.48 | 0.5283 | 0.6611 |

**Table 2:** Proboscidean DMTA summary values. A table containing the mean, median, range, and variation in *Asfc, epLsar*, and *Tfv* for each locality of *L. africana* and *M. colombi*.

| Locality | N | *Asfc* |  |  |  | *epLsar* |  |  |  | *Tfv* |  |  |  |
| --- | --- | --- | --- | --- | --- | --- | --- | --- | --- | --- | --- | --- | --- |
|  |  | Mean | Median | Range | SD (n-1) | Mean | Median | Range | SD (n-1) | Mean | Median | Range | SD (n-1) |
| Cypress Creek, TX<sup>1</sup> | 12 | 1.893 | 1.569 | 3.046 | 0.902 | 0.0039 | 0.0040 | 0.0052 | 0.0014 | 16056.38 | 16361.64 | 28621.02 | 8031.79 |
| Friesenhahn Cave, TX<sup>1</sup> | 31 | 2.921 | 2.592 | 6.014 | 1.53 | 0.0049 | 0.0048 | 0.0079 | 0.0021 | 12547.10 | 12462.43 | 12541.63 | 2665.38 |
| Ingleside, TX<sup>1</sup> | 26 | 2.285 | 2.063 | 5.481 | 1.27 | 0.0033 | 0.0030 | 0.0074 | 0.0019 | 10455.84 | 11304.11 | 17280.17 | 3710.62 |
| Leisey Shell Pit 1A, FL<sup>2</sup> | 17 | 1.906 | 1.817 | 3.23 | 0.865 | 0.0034 | 0.0034 | 0.0039 | 0.0012 | 15391.69 | 15803.26 | 25746.67 | 7169.71 |
| Punta Gorda, FL<sup>2</sup> | 11 | 2.565 | 2.442 | 3.282 | 1.07 | 0.0038 | 0.0035 | 0.0059 | 0.0015 | 24580.14 | 21479.48 | 55435.76 | 17042.09 |
| Tri-Britton, FL<sup>2</sup> | 5 | 3.932 | 3.317 | 4.874 | 2.21 | 0.0027 | 0.0027 | 0.0042 | 0.0017 | 11479.50 | 12011.90 | 8083.16 | 2945.24 |
| KNP, South Africa* | 11 | 3.835 | 3.397 | 6.53 | 1.93 | 0.0038 | 0.0039 | 0.0050 | 0.0014 | 13314.74 | 13673.70 | 9455.81 | 2470.48 |

<sup>1</sup>=Data are from Smith and DeSantis., 2018
<sup>2</sup>=Data are from Smith and DeSantis, 2020
\*Data are from this study, modern *Loxodonta africana* from Kruger National Park.

## Results

All DMTA values are reported in Tables 1 and 2. Significant differences were found for *Asfc* (chi-squared = 17.459, df = 6, p-value = 0.008), *epLsar* (Kruskal-Wallis chi-squared = 14.109, df = 6, p-value = 0.028), and *Tfv* (chi-squared = 16.987, df = 6, p-value = 0.009). A Dunn’s test found that median *Asfc* values from Leisey Shell Pit 1A (median = 1.817) differ significantly from those of the KNP sample (3.398; adjusted p = 0.045, Fig. 1); all other comparisons of the median values of DMTA parameters between fossil localities and the modern KNP sample were not significantly different (Table 2; Fig. 2-3). The overall pattern indicates a broad overlap in the dietary texture profiles across groups (Figs. 4 and 5), acknowledging the differences noted above.

**Fig. 1:**
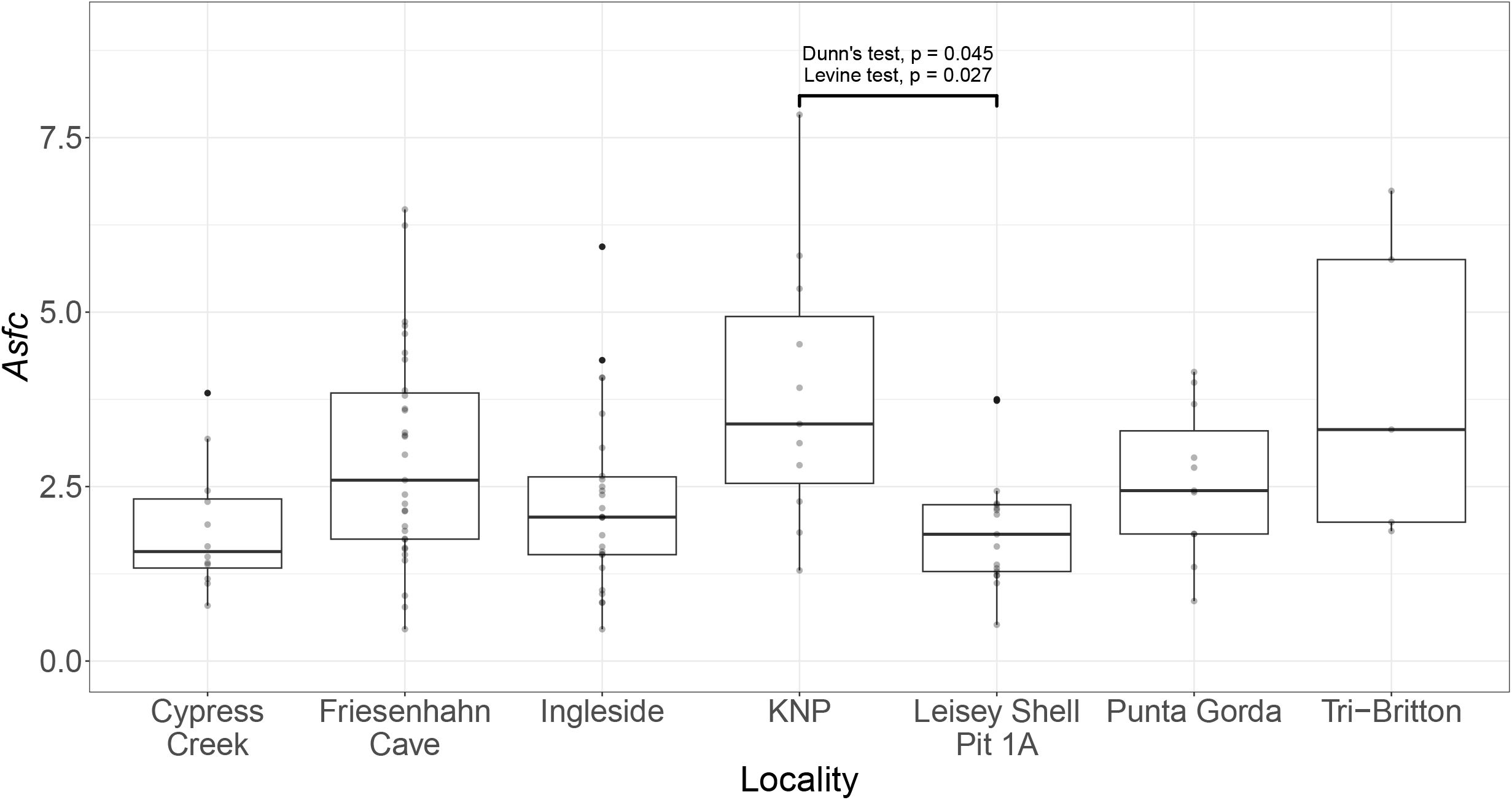
Mammoth and elephant complexity (*Asfc*) values across localities. A box plot comparing area-scale fractal complexity (*Asfc*) values of DMTA textures among the proboscidean specimens from each fossil and modern locality. Significant pairwise comparisons are indicated by the bracket, with the statistical test indicated.

**Fig. 2:**
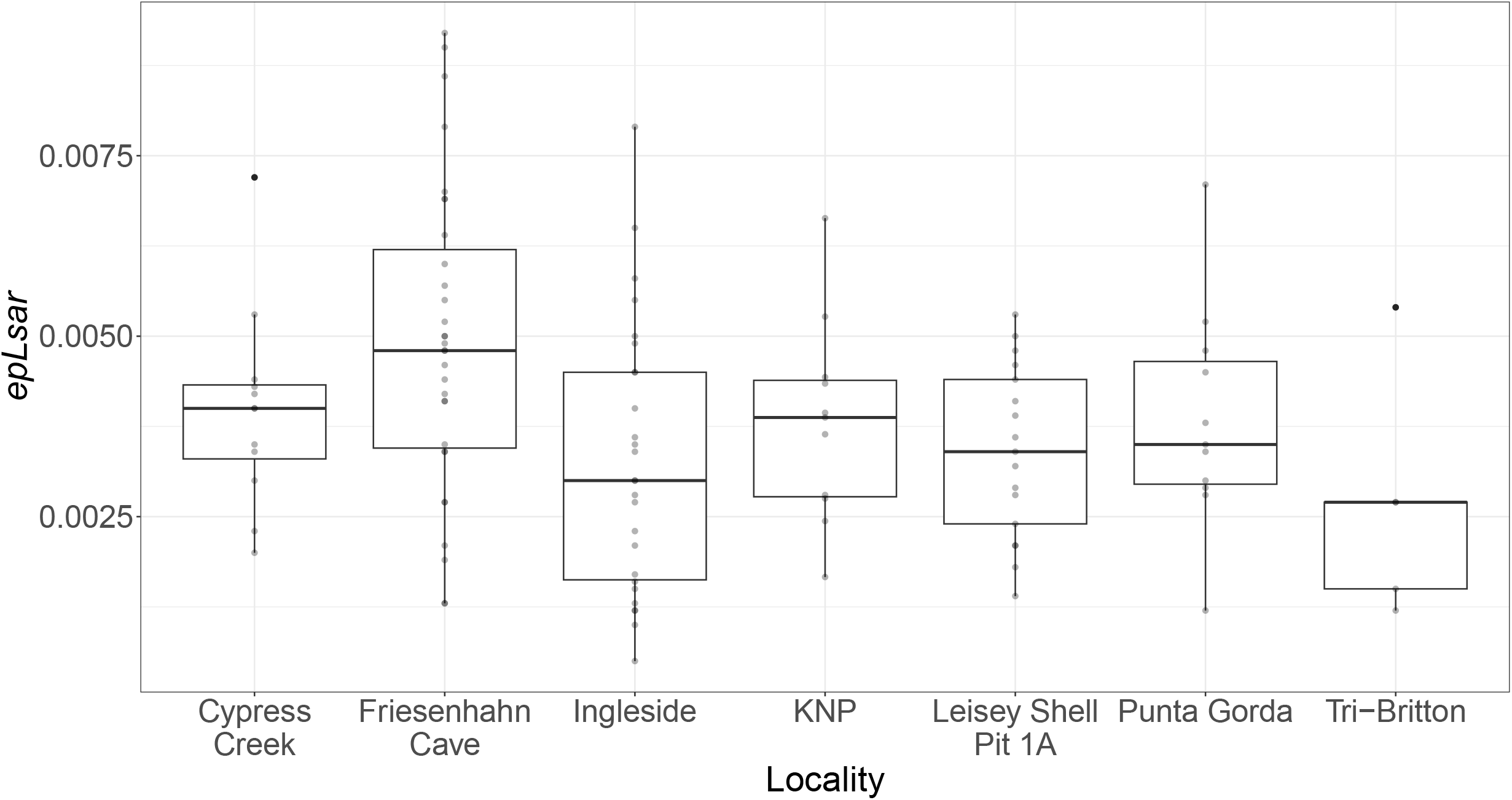
Mammoth and elephant anisotropy (*epLsar*) values across localities. A box plot of the distribution of *epLsar* values of DMTA textures among the proboscidean specimens from each fossil and modern locality.

**Fig. 3:**
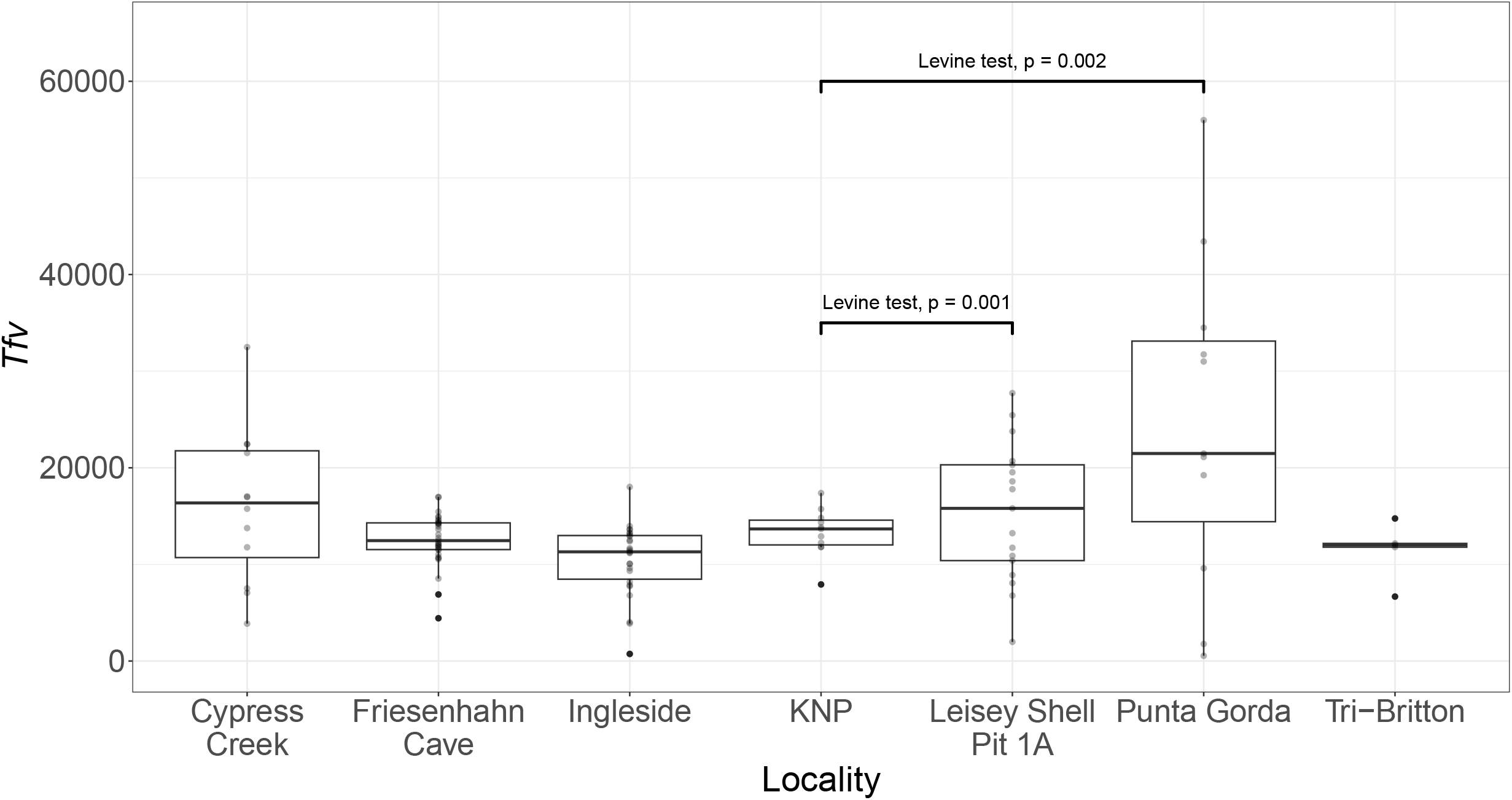
Mammoth and elephant textural fill volume (*Tfv)* values across localities. A box plot comparing *Tfv* values of DMTA textures among the proboscidean specimens from each fossil and modern locality. Significant pairwise comparisons are indicated by the bracket, with the statistical test indicated.

**Fig. 4:**
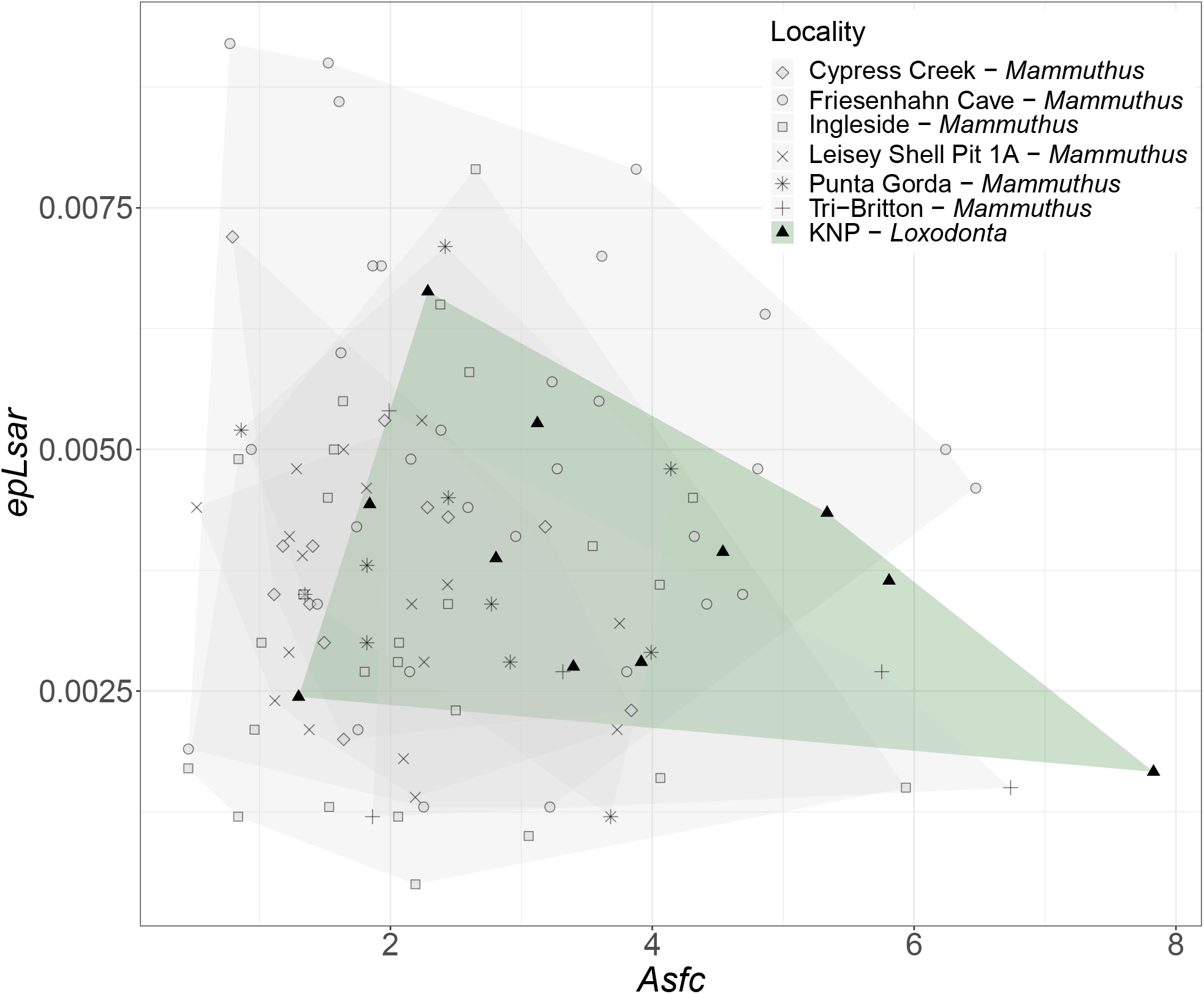
Biplot of anisotropy (*epLsar*) and complexity (*Asfc*). A scatterplot showing the relationship between anisotropy and complexity among all sampled individuals, with shading highlighting the space occupied by the microwear texture values of each locality. This plot visually compares the overlap of dietary texture patterns across extinct mammoth populations and the modern KNP elephants, allowing for assessment of dietary niche breadth.

**Fig. 5:**
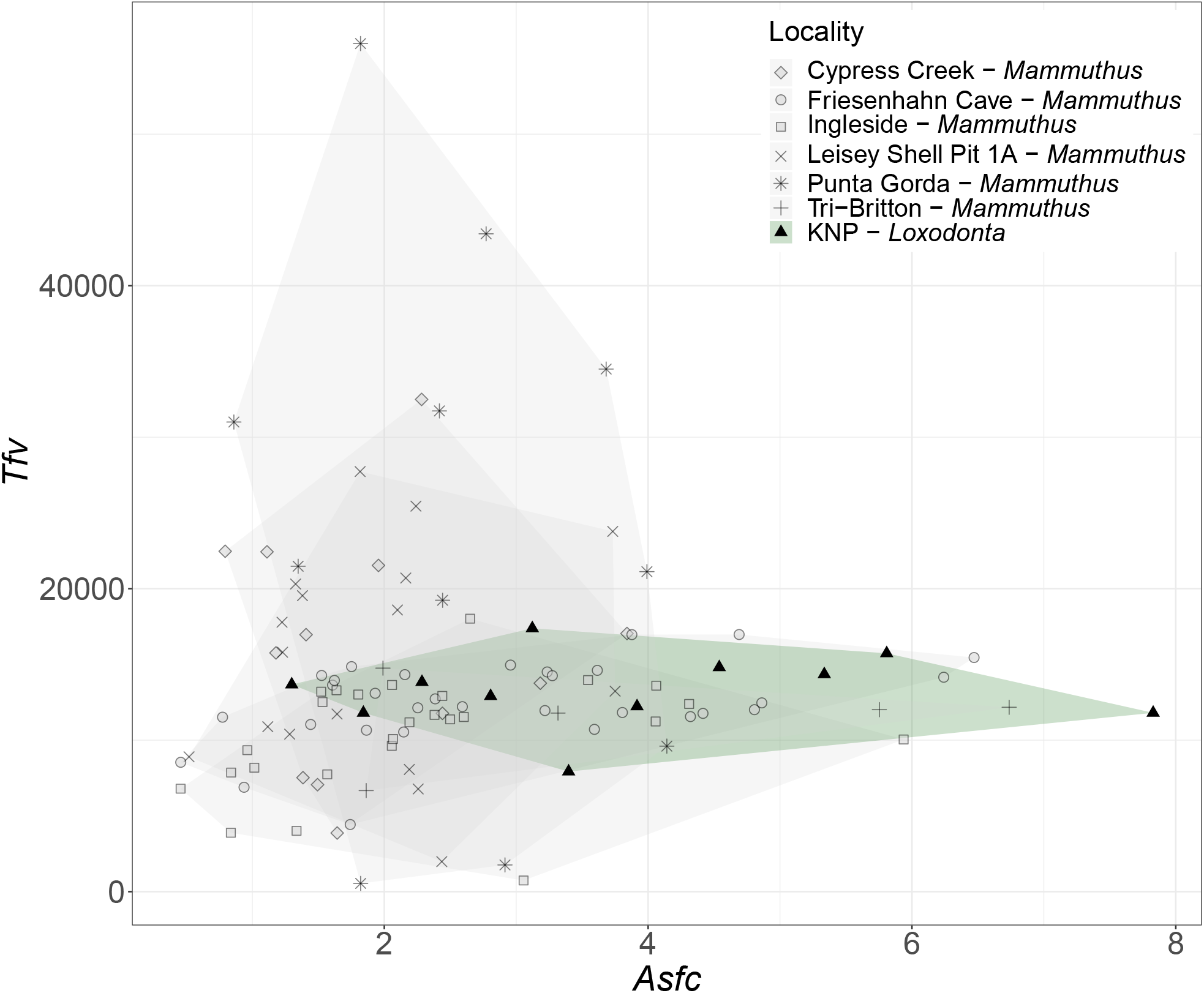
Biplot of textural fill volume (*Tfv*) and complexity (*Asfc*). Scatterplot of *Tfv* and *Asfc* values across all individuals, with shading highlighting the space occupied by the microwear texture values of each locality. This figure demonstrates how modern elephants and fossil mammoths compare in terms of surface relief and complexity.

Across our dataset, there are no significant differences in the variance of *epLsar* across localities (Levene test; df = 6, F = 1.2903, p = 0.268, Fig. 2), but there are significant differences in the variance of *Asfc* (df = 6, F = 2.3373, p = 0.037) and *Tfv* (df = 6, F = 11.865, p = 3.78×10^-10^). When we look at the pairwise comparisons of KNP’s variance in *Asfc* (3.72) and *Tfv* (6.10×10^6^) to the fossil localities using a post-hoc Levene test, we find few comparisons that are significantly different (α = 0.05). Leisey Shell Pit 1A has significantly different variance in *Asfc* (0.75, p = 0.027; Fig.1) and *Tfv* (5.14×10^7^, p = 0.001; Fig. 3) and Punta Gorda has significantly different variance in *Tfv* (2.90×10^8^, p = 0.002; Fig. 3). The overall pattern indicates broad similarities in the variance between the modern and fossil samples, acknowledging the differences above (Figs. 4 and 5). Notably, the variance in *Asfc* values measured from mammoths at Leisey Shell Pit 1A is lower than the KNP sample. Similarly, variance in *Tfv* values are lower at both Leisey Shell Pit 1A and Punta Gorda as compared to the KNP sample. The consistency in these DMTA variables further supports the interpretation that *Mammuthus columbi* processed foods with comparable mechanical properties similar to *Loxodonta africana* and likely relied on a mix of woody browse and grass in their diet.

## Discussion

Different dietary proxies capture dietary information over distinct temporal scales, with dental microwear reflecting short-term feeding behavior that occurred only days to weeks before death, sometimes referred to as the “Last Supper Effect” (Teaford and Oyen, 1989; Davis and Pineda-Munoz, 2016). In contrast, the individuals preserved at any given fossil locality can represent a range of temporal scales depending on the depositional context, from simultaneous mass death of multiple individuals to a highly time-averaged assemblage that has been accumulating over hundreds, thousands, or even millions of years (Kidwell, 1993; Kidwell, 2013; Fisher, 2018; Lundelius et al., 2019). A fundamental question in paleoecology is: To what extent are patterns measured from death assemblages reflective of real ecological phenomena? Can fossil assemblages be interpreted as a snapshot in time of populations, or does time averaging obscure the fidelity of the record?

Our constrained modern dataset from Kruger National Park provides a strong comparative baseline of what to expect from the microwear of a proboscidean population (Codron et al., 2006; Scott, 2012; Pineda-Munoz and Alroy, 2014), limiting the temporal snapshot to a few days within a given year and thus reflecting the past few weeks to months of these elephants’ diets. When compared to this reference sample, we find little evidence to support the hypothesis that time averaging inflates variation in DMTA parameters among mammoths. In fact, mammoths from Leisey Shell Pit 1A exhibit significantly lower variation in complexity (*Asfc*) than the KNP sample (Table 2, Fig. 1). These data further demonstrate the breadth of values that can be expected from proboscidean DMTA parameters, as the individuals from our sample of 11 African Bush elephants span the range previously reported across bovids of diverse feeding strategies (Scott, 2012).

Despite representing individuals sampled across long periods of time, fossil proboscidean assemblages do not exhibit inherently greater DMTA variability than the temporally and environmentally constrained population of *Loxodonta africana* from Kruger National Park. These findings challenge the assumption that time-averaged fossil datasets increase dietary variation and suggest that dental microwear may retain biologically meaningful dietary signals. This observation is consistent with stable isotope data, which show that high variability is not unique to fossil populations (DeSantis et al., 2017). For example, modern quokkas that died over the course of a decade from a ∼19 km^2^ island demonstrated nearly as much variability in oxygen isotope values as fossil assemblages potentially reflecting thousands of years of accumulation. Taken together, these results support interpretations of mammoths as flexible, generalist feeders and further validate the modern Kruger National Park elephants as an appropriate baseline for comparison (Smith et al., 2020).

The correspondence between fossil and modern microwear textures supports the use of *L. africana* as a valid analog for interpreting the dietary behavior of extinct proboscideans. As with all dietary proxies, we acknowledge the limitations in making direct comparisons between modern and fossil data. For example, the Kruger individuals were culled during the onset of the dry season, which is a transitional period for local vegetation. Given that these elephants consume more grass during the wet season (50% versus 40% in the dry season; Codron et al., 2006), it is possible that their DMTA values would have differed if sampled at another time of year. Additionally, grass consumption in the southern part of the park varies more seasonally (10% in the dry season, 50% in the wet season; Codron et al., 2006); thus, our sample may not capture the full spatial heterogeneity in DMTA values. Testing this remains difficult, as our sample was opportunistic and available as a result of controversial management activities that are no longer in practice. Still, the similarity in DMTA attributes between *L. africana* and fossil mammoths supports the interpretation that dietary flexibility as bulk mixed-feeders was a persistent feature of proboscidean ecology.

Our findings align with evidence from high-resolution fossil records showing that mammoths frequently responded to environmental variability through shifts in diet rather than through specialization. Seasonal and individual-level dietary changes documented in proboscidean tusks and other serial data reveal a capacity for short-term ecological adaptation, consistent with the microwear variability observed in both modern and extinct populations (Fisher, 2018; DeSantis et al., 2022). The lack of increased variation in mammoth microwear, despite the effects of time averaging, reinforces the conclusion that such flexibility was a fundamental aspect of proboscidean foraging behavior, much like *L. africana* today (Pineda-Munoz and Alroy, 2014). This result also aligns with broader research on the fidelity of time-averaged death assemblages, which shows that such assemblages can preserve biologically meaningful signals and reflect the ecological attributes of communities with surprising accuracy (Kidwell, 2013).

## Conclusions

This study demonstrates that the variation and breadth of dental microwear texture attributes in fossil mammoths are comparable to those observed in a temporally and environmentally constrained population of *Loxodonta africana*, a bulk mixed-feeder. Despite the long temporal span in fossil assemblages, mammoth DMTA data do not exhibit greater variability than modern elephants. These findings challenge the assumption that time averaging significantly inflates dietary variation in fossil taxa and instead suggest that dental microwear preserves ecologically meaningful signals even across long timescales. These results support the interpretations of mammoths and modern elephants as dietary generalists with a high degree of ecological flexibility. This dietary plasticity has been a persistent feature of proboscidean ecology. Further, these comparisons affirm the value of *L. africana* as a modern analog for extinct mammoths and underscores the utility of DMTA as a reliable proxy for reconstructing short-term dietary behavior in modern and fossil assemblages.

By establishing a baseline of dental microwear variation in a well-documented modern population, this study provides a framework for more accurately interpreting dietary ecology in extinct megafauna. These findings reinforce the ecological fidelity of time-averaged fossil assemblages and support the broader application of high-resolution proxies in paleoecological reconstructions. Future research should expand this comparative approach to include additional modern populations, explore seasonal dietary shifts, and integrate multi-proxy methods such as isotopic and mesowear analyses. Such work will continue to refine our understanding of both extinct and extant herbivore ecology and enhance the interpretive power of the fossil record.

## Acknowledgements

We would like to express our sincere gratitude to all those who have contributed to the success of this project. A special thanks to Vanderbilt University (Department of Biological Sciences; Evolutionary Studies Initiative; Department of Earth & Environmental Studies) and the Illinois State Museum for access to labs, equipment, and collections, as well as Elsa Mueller-Filipes for assistance in data collection and Aditya Kurre for assistance with confocal microscopy and data interpretation.

## References

Abraham, J. O., Rowan, J., O’Brien, K., Sokolowski, K. G., & Faith, J. T. (2024). Environmental context shapes the relationship between grass consumption and body size in African herbivore communities. Ecology and Evolution, 14(2), e11050. 10.1002/ece3.11050

Behrensmeyer, A. K. (1982). Quaternary palaeoecology. By H.J.B. Birks and Hilary H. Birks. Baltimore: University Park Press. 1980. 289 pp., figures, tables, bibliographies, index. $84.50 (cloth). American Journal of Physical Anthropology, 58(1), 119–120. 10.1002/ajpa.1330580119

Codron, J., Lee-Thorp, J. A., Sponheimer, M., Codron, D., Grant, R. C., & de Ruiter, D. J. (2006). Elephant (Loxodonta africana) Diets in Kruger National Park, South Africa: Spatial and Landscape Differences. Journal of Mammalogy, 87(1), 27–34. 10.1644/05-MAMM-A-017R1.1

Davis, M., & Pineda-Munoz, S. (2016). The temporal scale of diet and dietary proxies. Ecology and Evolution, 6(6), 1883–1897. 10.1002/ece3.2054

DeSantis, L. R., Schubert, B. W., Schmitt-Linville, E., Ungar, P. S., Donohue, S. L., & Haupt, R. J. (2015). Dental microwear textures of carnivorans from the La Brea Tar Pits, California and potential extinction implications. Contributions in Science, Los Angeles County Museum of Natural History, 42, 37–52.

DeSantis, L. R. G. (2016). Dental microwear textures: Reconstructing diets of fossil mammals. Surface Topography: Metrology and Properties, 4(2), 023002. 10.1088/2051-672X/4/2/023002

DeSantis, L. R. G., Crites, J. M., Feranec, R. S., Fox-Dobbs, K., Farrell, A. B., Harris, J. M., Takeuchi, G. T., & Cerling, T. E. (2019). Causes and Consequences of Pleistocene Megafaunal Extinctions as Revealed from Rancho La Brea Mammals. Current Biology, 29(15), 2488-2495.e2. 10.1016/j.cub.2019.06.059

DeSantis, L. R. G., Field, J. H., Wroe, S., & Dodson, J. R. (2017). Dietary responses of Sahul (Pleistocene Australia–New Guinea) megafauna to climate and environmental change. Paleobiology, 43(2), 181–195. 10.1017/pab.2016.50

DeSantis, L. R., Pardi, M. I., Du, A., Greshko, M. A., Yann, L. T., Hulbert Jr, R. C., & Louys, J. (2022). Global long-term stability of individual dietary specialization in herbivorous mammals. Proceedings of the Royal Society B, 289(1968), 20211839. 10.1098/rspb.2021.1839

DeSantis, L. R. G., Sharp, A. C., Schubert, B. W., Colbert, M. W., Wallace, S. C., & Grine, F. E. (2020). Clarifying relationships between cranial form and function in tapirs, with implications for the dietary ecology of early hominins. Scientific Reports, 10(1), 8809. 10.1038/s41598-020-65586-w

Feranec, R. S., & MacFadden, B. J. (2000). Evolution of the grazing niche in Pleistocene mammals from Florida: Evidence from stable isotopes. Palaeogeography, Palaeoclimatology, Palaeoecology, 162(1), 155–169. 10.1016/S0031-0182(00)00110-3

Fisher, D. C. (2018). Paleobiology of Pleistocene Proboscideans. Annual Review of Earth and Planetary Sciences, 46(Volume 46, 2018), 229–260. 10.1146/annurev-earth-060115-012437

Green, J. L., DeSantis, L. R. G., & Smith, G. J. (2017). Regional variation in the browsing diet of Pleistocene Mammut americanum (Mammalia, Proboscidea) as recorded by dental microwear textures. Palaeogeography, Palaeoclimatology, Palaeoecology, 487, 59–70. 10.1016/j.palaeo.2017.08.019

Grine, F. E., & Kay, R. F. (1988). Early hominid diets from quantitative image analysis of dental microwear. Nature, 333(6175), 765–768. 10.1038/333765a0

Haupt, R. J., DeSantis, L. R. G., Green, J. L., & Ungar, P. S. (2013). Dental microwear texture as a proxy for diet in xenarthrans. Journal of Mammalogy, 94(4), 856–866. 10.1644/12-MAMM-A-204.1

Hedberg, C., & DeSantis, L. R. G. (2017). Dental microwear texture analysis of extant koalas: Clarifying causal agents of microwear. Journal of Zoology, 301(3), 206–214. 10.1111/jzo.12413

Jones, D. B., & Desantis, L. R. G. (2016). Dietary Ecology of the Extinct Cave Bear: Evidence of Omnivory as Inferred from Dental Microwear Textures. Acta Palaeontologica Polonica, 61(4), 735–741. 10.4202/app.00253.2016

Jones, D. B., & Desantis, L. R. G. (2017). Dietary ecology of ungulates from the La Brea tar pits in southern California: A multi-proxy approach. Palaeogeography, Palaeoclimatology, Palaeoecology, 466, 110– 127. 10.1016/j.palaeo.2016.11.019

Kamga, S. M., Tamungang, S. A., Awa, T., Ewome, F. L., Motombi, F. N., Hořák, D., & Riegert, J. (2022). The Importance of Forest Elephants for Vegetation Structure Modification and Its Influence on the Bird Community of a Mid-Elevation Forest on Mount Cameroon, West-Central Africa. Diversity, 14(3), Article 3. 10.3390/d14030227

Kidwell, S. M. (1993). Patterns of time-averaging in the shallow marine fossil record. Short Courses in Paleontology, 6, 275–300. 10.1017/S247526300000115X

Kidwell, S. M. (2013). Time-averaging and fidelity of modern death assemblages: Building a taphonomic foundation for conservation palaeobiology. Palaeontology, 56(3), 487–522. 10.1111/pala.12042

Kidwell, S. M., & Tomasovych, A. (2013). Implications of Time-Averaged Death Assemblages for Ecology and Conservation Biology. Annual Review of Ecology, Evolution, and Systematics, 44(Volume 44, 2013), 539–563. 10.1146/annurev-ecolsys-110512-135838

Koch, P. L., Diffenbaugh, N. S., & Hoppe, K. A. (2004). The effects of late Quaternary climate and pCO2 change on C4 plant abundance in the south-central United States. Palaeogeography, Palaeoclimatology, Palaeoecology, 207(3), 331–357. 10.1016/j.palaeo.2003.09.034

Lundelius, E. L., Thies, K. J., Graham, R. W., Bell, C. J., Smith, G. J., & DeSantis, L. R. G. (2019). Proboscidea from the Big Cypress Creek fauna, Deweyville Formation, Harris County, Texas. Quaternary International, 530–531, 59–68. 10.1016/j.quaint.2019.11.018

Owen-Smith, N. (1987). Pleistocene extinctions: The pivotal role of megaherbivores. Paleobiology, 13(3), 351–362. 10.1017/S0094837300008927

Pineda-Munoz, S., & Alroy, J. (2014). Dietary characterization of terrestrial mammals. Proceedings of the Royal Society B: Biological Sciences, 281(1789), 20141173. 10.1098/rspb.2014.1173

Poulsen, J. R., Rosin, C., Meier, A., Mills, E., Nuñez, C. L., Koerner, S. E., Blanchard, E., Callejas, J., Moore, S., & Sowers, M. (2018). Ecological consequences of forest elephant declines for Afrotropical forests. Conservation Biology: The Journal of the Society for Conservation Biology, 32(3), 559–567. 10.1111/cobi.13035

Prideaux, G. J., Ayliffe, L. K., DeSantis, L. R. G., Schubert, B. W., Murray, P. F., Gagan, M. K., & Cerling, T. E. (2009). Extinction implications of a chenopod browse diet for a giant Pleistocene kangaroo. Proceedings of the National Academy of Sciences, 106(28), 11646–11650. 10.1073/pnas.0900956106

Rivals, F., Solounias, N., & Mihlbachler, M. C. (2007). Evidence for geographic variation in the diets of late Pleistocene and early Holocene Bison in North America, and differences from the diets of recent Bison. Quaternary Research, 68(3), 338–346. 10.1016/j.yqres.2007.07.012

Scott, J. R. (2012). Dental microwear texture analysis of extant African Bovidae. 76(2), 157–174. 10.1515/mammalia-2011-0083

Scott, R. S., Teaford, M. F., & Ungar, P. S. (2012). Dental microwear texture and anthropoid diets. American Journal of Physical Anthropology, 147(4), 551–579. 10.1002/ajpa.22007

Scott, R. S., Ungar, P. S., Bergstrom, T. S., Brown, C. A., Grine, F. E., Teaford, M. F., & Walker, A. (2005). Dental microwear texture analysis shows within-species diet variability in fossil hominins. Nature, 436(7051), 693–695. 10.1038/nature03822

Smith, G. J., & DeSantis, L. R. G. (2018). Dietary ecology of Pleistocene mammoths and mastodons as inferred from dental microwear textures. Palaeogeography, Palaeoclimatology, Palaeoecology, 492, 10– 25. 10.1016/j.palaeo.2017.11.024

Smith, G. J., & DeSantis, L. R. G. (2020). Extinction of North American Cuvieronius (Mammalia: Proboscidea: Gomphotheriidae) driven by dietary resource competition with sympatric mammoths and mastodons. Paleobiology, 46(1), 41–57. 10.1017/pab.2020.7

Teaford, M. F., & Oyen, O. J. (1989). In vivo and in vitro turnover in dental microwear. American Journal of Physical Anthropology, 80(4), 447–460. 10.1002/ajpa.1330800405

Ungar, P. S., Brown, C. A., Bergstrom, T. S., & Walker, A. (2003). Quantification of dental microwear by tandem scanning confocal microscopy and scale-sensitive fractal analyses. SCANNING, 25(4), 185–193.

van Aarde, R., Whyte, I., & Pimm, S. (1999). Culling and the dynamics of the Kruger National Park African elephant population. ANIMAL CONSERVATION, 2(4), 287–294. 10.1017/S1367943099000621

Walker, A., Hoeck, H. N., & Perez, L. (1978). Microwear of Mammalian Teeth as an Indicator of Diet. Science, 201(4359), 908–910. 10.1126/science.684415

White, J. M., DeSantis, L. R. G., Evans, A. R., Wilson, L. A. B., & McCurry, M. R. (2021). A panda-like diprotodontid? Assessing the diet of Hulitherium tomasettii using dental complexity (Orientation Patch Count Rotated) and dental microwear texture analysis. Palaeogeography, Palaeoclimatology, Palaeoecology, 583, 110675. 10.1016/j.palaeo.2021.110675

Yann, L. T., & DeSantis, L. R. G. (2014). Effects of Pleistocene climates on local environments and dietary behavior of mammals in Florida. Palaeogeography, Palaeoclimatology, Palaeoecology, 414, 370–381. 10.1016/j.palaeo.2014.09.020

